# Cyanochelin B: A siderophore produced by cyanobacterium *Leptolyngbya* sp. NIES-3755 with photolytic properties that negate iron monopolization in the UV-light

**DOI:** 10.1101/2025.01.08.631963

**Authors:** Berness P. Falcao, Viviana Di Matteo, Pavel Hrouzek, Lenka Štenclová, Petra Urajová, Jan Mareš, Jan Kuta, José Alberto Martínez Yerena, Eliška Kozlíková-Zapomělová, Germana Esposito, Alfonso Mangoni, Valeria Costantino, Tomáš Galica

## Abstract

Siderophores are low-molecular-weight compounds excreted by microorganisms to facilitate iron uptake in times of its unavailability. Microbes may produce siderophores to monopolize iron and achieve competitive exclusion of other strains. Alternatively, siderophores may be exchanged for other substrates in mutualistic relationships. Siderophores that employ *β*-hydroxy-aspartate (*β*-OH-Asp) for iron chelation were shown to undergo UV-mediated photolytic cleavage with simultaneous reduction of Fe^3+^ to Fe^2+^. Photolytic siderophores can mediate algal-bacterial mutualism, where the bacteria provide iron in exchange for dissolved organic carbon.

We use an interdisciplinary strategy to provide a complex characterization of cyanochelin B, a photolytic β-OH-Asp-containing siderophore produced by filamentous cyanobacterium *Leptolyngbya* sp. NIES-3755. A combination of nuclear magnetic resonance, high resolution mass spectrometry and bioinformatic analyses complemented with Marfey’s and Murata’s methods yielded the structure of cyanochelin B with the configuration of its stereocenters. Cyanochelin B-iron complexes exposed to UV-light photolyze within minutes (t_1/2_ = 8.9 min; ∼3.5 uE UV-A) and release reduced Fe^2+^. We have co-cultured *Leptolyngbya* together with *Synechocystis* PCC6803 as a reporter strain lacking siderophore production. The cultivation setup was based on membrane-separated compartments accommodating individual strains and employed alginate-embedded FeCl_3_ to simulate poorly accessible precipitated iron. Our results demonstrate that in the absence of UV-light cyanochelin B can monopolize iron in favor of *Leptolyngbya.* However, UV-light eliminates any monopolization of iron and makes it available to competing organisms. Finally, we report the isolation of novel cyanochelin B-producing strains of *Phormidesmis* from field material and discuss the phenomenon of photolytic siderophores in a broader context.

**Importance:** Iron is an essential micronutrient that is required by all living organisms as a cofactor of indispensable enzymes. Due to its specific properties it is mostly precipitated and biologically unavailable. To facilitate iron uptake, microbes produce siderophores - low-molecular-weight compounds that bind iron. Siderophores are mediators of microbial interactions and facilitate competitive exclusion of non-compatible strains or foster mutualistic partners and cheater strains. We report a full structural elucidation of cyanochelin B, a photolytic cyanobacterial siderophore that contains β-OH-Asp. Our co-culture experiments show that cyanochelin B may monopolize iron to its producer or make it accessible to other strains depending on the presence of UV light. Moreover, our data suggest that the benefits from production of photolytic siderophores are not restricted to symbiotic partners of the producer but rather available to the whole irradiated community. Of known siderophores, 17.5% contain the photoreactive β-OH-Asp and can function in a similar way.

## Introduction

Cyanobacteria are photoautotrophic prokaryotes that use inorganic carbon and solar energy (Whitton et al 2012). In environmental conditions that do not favor land plants or macroscopic algae, they often serve as key primary producers of a given habitat and provide the surrounding microbial community with fixed organic carbon (Campbell 1979, Oren et al 2017). Although cyanobacteria and heterotrophic bacteria use complementary sources of carbon and energy, they often compete for many other nutrients. Iron is an essential micronutrient, vital to the enzymes of the respiratory chain and the photosynthetic machinery, and hence required by both phototrophs and heterotrophs alike (Kramer et al 2019, Arstol et al 2019). While iron is the fourth most abundant element in Earth’s crust, its complicated chemistry makes it often inaccessible and efficiently the limiting element, such as in the ocean gyres (Sandy and Butler 2009, Martin et al 1994).

During evolution, cyanobacteria and other bacteria, adopted various strategies to cope with iron limitation, such as storing iron reserves or producing siderophores - iron-chelating compounds that facilitate iron uptake (Mazzotta et al 2020, Arstol et al 2019, Keren et al 2004). In addition to iron-chelating residues, such as catecholate, hydroxamate or carboxylate, siderophores can contain additional structural features, e.g. modified amino or fatty acids, to fine-tune their functional properties. The combination of the residues employed results in over 700 unique structures described thus far (Siderite database, He et al 2024). The repertoire of known cyanobacterial siderophores, however, counts only 19 structures in total - schizokinen, 6 variants of synechobactin, 3 variants of anachelin, 6 variants of cyanochelins and 3 leptochelins (Avalon 2024 et al, Galica et al 2021, Arstol et al 2019).

Siderophores are released into the environment and once iron is bound, the iron-siderophore complexes are taken up and processed by a multi-step transport machinery. Siderophore-producing bacteria possess importers that efficiently recognize and retrieve iron bound by their own siderophore and thus gain access to additional sources of iron. Cells lacking a compatible siderophore importer are locked away from the siderophore bound iron and in time may be outcompeted (Kramer et al 2019). However, many microbes that do not produce siderophores can purposefully take up the siderophores produced by others and thus avoid the metabolic costs of siderophore production while attaining its benefit (Kramer et al 2019, Tostado-Islas et al 2021). In extreme cases, such microbes depend on specific siderophores produced by other species as their sole mean of iron uptake (D’Onofrio et al 2010). Thus, siderophores limit the access to iron exclusively for the cells with appropriate transporters, although they may not produce the siderophores themselves.

Interestingly, many siderophores that bind iron via β-hydroxy-aspartates (β-OH-Asp) lyse upon exposure to UV light and during the process reduce the bound Fe^3+^ to Fe^2+^ (Butler et al 2021, Kreutzer et al 2012, Hardy and Butler 2019). The latter is more soluble in neutral pH and is easily and non-exclusively taken up by cells. Photolytically reduced iron is thought to be a service offered by bacteria to siderophore non-producing eukaryotic algae in return for provided dissolved organic carbon (Barbeau et al 2001, Jiang et al 2024). However, this is unlikely to be a relevant explanation for siderophore-producing photoautotrophic cyanobacteria. Apparently, siderophores can be a focal point of complex microbial interactions in iron limited communities.

Previously, we described a novel class of cyanobacterial siderophores named cyanochelins (Galica et al 2021). Cyanochelins feature two β-OH-Asp for iron chelation, can perform a photolytic reduction of ferric iron, and according to our bioinformatic survey can be expected to be widely distributed. Recently we have detected cyanochelin B in a field-obtained microbial community that was experimentally starved for iron. In the present paper, we report full structural elucidation of cyanochelin B and investigate the effect that UV-dependent photolysis of iron-cyanochelin complexes has on the distribution of iron in a coculture system.

## Results

### Structure of Cyanochelin B

Cyanochelin B (2 mg) at 99% purity was obtained using high-performace liquid chromatography (HPLC) fractionation of refined biomass extract (for details see methods section) of iron-starved *Leptolyngbya* sp. NIES 3755. The purified compound was subjected to a full set of 1D and 2D homo- and hetero-nuclear magnetic resonance (NMR) experiments complemented by high resolution mass spectrometry analysis (HRMS). Evaluation of cyanochelin B structure was further supported by bioinformatic analysis of its putative BGC and Marfey’s and Murata’s methods for determination of stereochemistry (Marfey et al 1984, Matsumori et al 1999, Galica et al 2021).

The HRMS spectrum of cyanochelin B (Fig. S1, Table S1) showed a molecular ion peak [M+H]^+^ at *m/z* 1026.5111 corresponding to the molecular formula C_47_H_75_N_7_O_16_S (Fig. S1). The analysis of the MS/MS spectrum displayed a characteristic fragment ion at *m/z* 786.2613, which corresponds to a loss of the N-terminal hydroxylated hydrocarbon chain cleaved between C30 and C32, yielding a peptide core of the siderophore (M-FA, Fig. S1). Subsequently, the peptide core yields an MS/MS spectrum corresponding to the sequential losses of individual amino acid/residues starting from the *C*-terminus and forming fragments at 725.2580 (b6-FA, loss of ethanolamine), 594.1889 (b5-FA, loss of β-OH-Asp), 507.1543 (b4-FA, loss of Ser), 376.1325 (b3-FA, loss of β-OH-Asp), 319.1107 (b2-FA, loss of Gly) and 144.0474 (a1-FA. loss of Phe) and was in congruence with the specificity and order of A-domains found in the NRPS-PKS biosynthetic gene cluster (BGC) located on plasmid 2 of *Leptolyngbya* sp. NIES-3755 (GenBank: AP017310.1, for biosynthesis and MS fragmentation see supplementary data, Table S1, Table S2, Fig. S1 and Fig. S2). The peptide substructure was elucidated by a full set of NMR data (Table S3), i.e., by the presence of five distinct amide-NH signals (δ_H_-DMSO of NH-17: 7.65 ppm, NH-15: 8.27 ppm, NH-11: 8.07 ppm, NH-8: 7.83 ppm, NH-4: 7.93), four signals related to α-amino acid protons (δ_H_-DMSO of H-4: 4.69 ppm, H-8: 4.45 ppm, H-11: 4.76 ppm, H-17: 4.60 ppm), two signals related to methylene protons attributable to a glycine residue (δ_H_-DMSO of H-15a: 3,89 ppm and H-15b: 3,80 ppm). Two signals related to methine protons binding oxygen, C-5 and C-12, resonating at δ_H_-DMSO 4.60 ppm and 4.51 ppm, respectively, confirmed the localization of hydroxyl groups on β-carbon of aspartate residues. Finally, the presence of amide signal (δ_H_-DMSO of NH-2: 7.68 ppm) and COSY and HMBC correlations suggest that the C-terminal residue is ethanolamine, presumably derived from a Gly reduced to a primary alcohol by a thioesterase module with reductive function found to be encoded alongside a Gly-specific NRPS module in likely the last gene of the deduced BGC (BAU16060; for details see Fig. S2).

The ¹H-NMR signals indicated the presence of a saturated hydrocarbon chain. Typical methyl triplet at 0.86 ppm (C-47) is correlated in the COSY spectrum to the methylene resonating at 1.26 ppm (C-46) and is followed by a 14-methylene chain (δ_H_ from 1.23 to 1.51 ppm, δ_C_ from 22.1 to 31.3 ppm). This 15-carbon chain is attached to the methine carbon binding oxygen (δ_H_-32: 3.47 ppm, δ_C_-32: 76 ppm), which is in turn connected to the quaternary carbon (δ_C_-30: 76.8 ppm) that binds an oxygen and a methyl group C-31 (δ_H_-31: 1.34 ppm, δ_C_-31: 24.0 ppm), as is confirmed by the HMBC correlation shown in Fig. 1. The quaternary carbon is linked to a methyl-thiazoline unit, as evidenced by HMBC correlation C-31/C-29 and C-27/C-29. The presence of a methyl-thiazoline residue is expected due to the presence of a predicted cysteine-specific NRPS module with a cyclization domain and a methyltransferase domain (Fig. 1. and Fig. S2) and is congruent with the obtained HMBC and NOESY correlations (Table S3, Fig. S7 and Fig. S9) and due to the presence of sulfur also with the isotopic pattern of fragment 144.0474 observed by HRMS.

**FIG 1.**
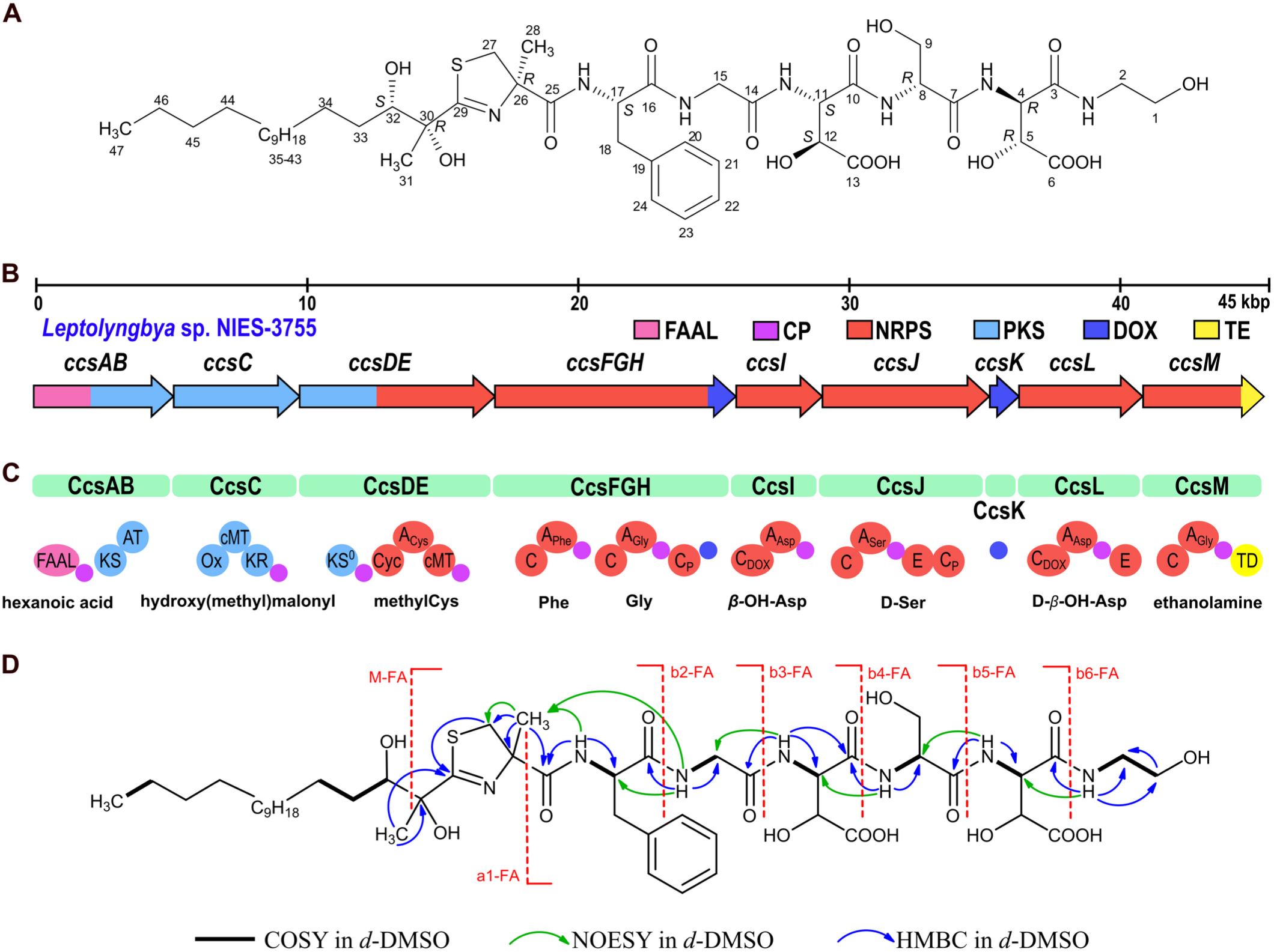
**(A)** Structure and stereochemistry of cyanochelin B. **(B)** Architecture of the gene cluster encoding hybrid PKS/NRPS biosynthesis of cyanochelin B (BGC). Genes are color-coded according to the type of biosynthetic step: FAAL - fatty acyl-AMP ligase, pink; CP - carrier protein, purple; NRPS - non-ribosomal peptide synthetase, red; PKS - polyketide synthase, light blue; DOX - aspartate β-hydroxylase, blue; TE - thioesterase, yellow. **(C)** Modules of the PKS/NRPS encoded by the cluster: KS, β-ketoacyl synthase; KS^0^, non-elongating β-ketoacyl synthase; AT, acyltransferase; Ox, flavin-containing monooxygenase; cMT, C-methyltransferase domain; KR, β-ketoacyl reductase; Cyc, cyclization (thiazole-forming) condensation domain; C, condensation domain; A, adenylation domain; E, epimerase domain; TD, thioester reductase domain. Subscript specifies the type of C domains or the specificity of A-domains. **(D)** Most relevant 2D NMR correlations and MS fragments of cyanochelin B.

Cyanochelin B thus consists of [acylated thiazoline^1^, Phe^2^, Gly^3^, β-OH-Asp^4^, Ser^5^, β-OH-Asp^6^, ethanolamine^7^] and contains 9 stereocenters (Fig. 1). The Marfey’s method was applied to determine the absolute configuration of the α-stereocenters of amino acids and suggested the presence of L-Phe (*S*-Phe), D-Ser (*R*-Ser), and both the L-(2*S*,3*S*) and D-(2*R*,3*R*) forms of *threo* β-OH-Asp (Fig. S12 and S13). The observation aligns with the presence of epimerase domains in NRPS modules responsible for incorporating Ser^5^ and β-OH-Asp^6^. The placement of L- and D-enantiomers of *threo*-β-OH-Asp within cyanochelin B was not apparent at first. However, a careful examination of the NMR data of β-OH-Asp^6^ moiety revealed that the proton H-4, resonating as a double doublet at δ 4.69, is coupled to H-5 with a ^3^J_H,H_ coupling constant of 2.3 Hz (Table S3, Fig. S10). Furthermore, a comparison of the multiplicity of H-4 and H-5 in the normal ¹H NMR spectrum and in the 1D sections of the HMBC spectrum taken at δ 173.3 (C-6) and δ 168.8 (C-3), respectively, showed that the ^3^*J*_C,H_ coupling constants between H-4 and C-6 and between H-5 and C-3 are both small (Fig. S10). The combined data enabled the application of the Murata’s method, establishing the relative configuration of β-OH-Asp^6^ as *threo*. Similar results were obtained for β-OH-Asp^4^, which also has a *threo* relative configuration. (Fig. S11). Bioinformatic analysis of the biosynthetic gene cluster (BGC) suggests the same configuration. Reitz and colleagues investigated the stereochemistry of β-OH-Asp in siderophores and reported that with one exception, all β-OH-Asp in siderophores are either D-*threo* or *L-threo* (Reitz et al 2020). According to their study, the NRPS-fused β-hydroxylases should produce *S* while the self-standing enzymes should produce *R* configuration at the β-carbon. The NRPS module that should incorporate β-OH-Asp at position 4 lacks an epimerase (CcsI) and is preceded by NRPS-fused β-hydroxylase (CcsFGH), hence the amino acid at position 4 should be L-*threo*-β-OH-Asp (11*S*,12*S*). The NRPS module responsible for the incorporation of amino acid at position 6, CcsL (BAU16061), is Asp-specific, contains an epimerase domain and is preceded by a self-standing β-hydroxylase (CcsK) and should produce D-*threo* β-OH-Asp (4*R*,5*R*).

As for the configuration at C-26 of thiazoline, a methylcysteine is formed from the opening of the thiazoline ring during the hydrolysis of cyanochelin B. Using the Marfey’s method, it was determined that this compound is L-methylcysteine (*R*-methylcysteine). Although we did not have an authentic sample of either D- or L-methylcysteine, we found relevant literature that provided useful information. According to Carmeli et al. (1993), the L-methylcysteine-L-DAA derivative exhibits longer retention times than the D-methylcysteine-L-DAA derivative. Therefore, the L-DAA derivative, showing a longer retention time (Figure S12), must be L-methylcysteine-L-DAA. In comparison, the D-DAA derivative with a shorter retention time must be L-methylcysteine-D-DAA, enantiomeric with D-methylcysteine-L-DAA.

The ketoreductase domain of CcsC (BAU16015) is expected to perform a stereospecific reduction of the β-keto group at C-32 (Keatinge-Clay 2007). Due to the absence of conserved LDD and HXXY motifs, the reduction of the oxo group at C32 yields a hydroxyl in the *S* configuration at the respective position (Keatinge-Clay 2007). The NMR measurements suggest the relative configuration at C-30/C-32 to be *threo* (either 30*S*,32*R* or 30*R*,32*S*) because the multiplicity of H-32, resonating as a broad doublet (J = 9.9 Hz) aligns with that reported in the literature for the *threo* stereoisomers of methyl 2,3-dihydroxy-2-methyloctanoate (Chatzimpaloglou et al 2014; Chatzimpaloglou et al 2012). Taking together specificity analysis of CcsC and the NMR data we conclude that the configuration at C-30/C-32 of cyanochelin B is a *threo* 30*R* and 32*S*.

### Kinetics of photolysis and structure of photolytic fragments

Siderophores containing a β-OH-Asp in their structure can reduce the bound ferric iron if exposed to UV-light with simultaneous photolytic cleavage of the siderophore molecule. To understand how cyanochelin B functions in nature, as well as in our laboratory experiments, we have determined cyanochelin B half-life at different UV-light intensities.

Cyanochelin B showed no or very mild decomposition in absence of UV light or ferric iron. In presence of UV light, the iron-siderophore complexes underwent a photolytic reaction with 0-order kinetics and half-life 8.9 minutes in the highest UV irradiation tested (Fig. 2.) and 24.1 minutes in the light conditions that were later used in the co-cultivation experiments (½ UV intensity, including lid of the well plate).

**Fig. 2.**
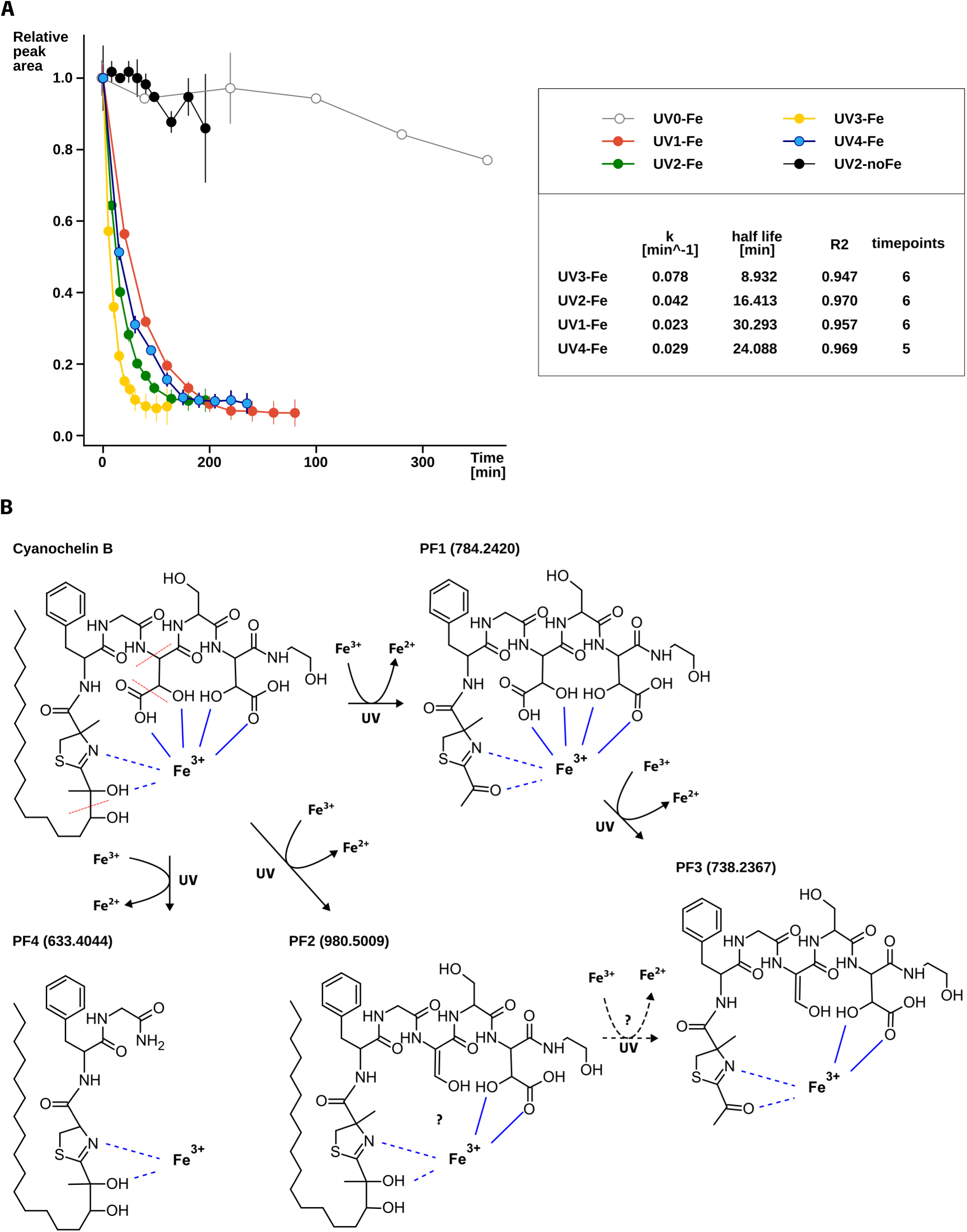
Photolysis of cyanochelin B. **A)** Kinetics of photolysis of cyanochelin B depicted as decrease in relative peak area calculated from HPLC-HRMS analyses over time (see methods). Legend and table with the calculated reaction rates/half-lives and R2 score from linear regression of log-transformed data are in the inset. Time points at which the relative peak area is close to zero were omitted from the regression analysis. **B)** Scheme of possible photolytic reactions of cyanochelin B and resulting photolytic fragments.

Further HPLC-HRMS analysis revealed several new compounds - candidate photolytic fragments (PF1-PF4, Fig. 2, Fig. S14-S17). Their MS/MS spectra contained signature ions confirming that the compounds originate from cyanochelin B (Fig. S14-S17) and provided information to identify the site of photolytic cleavage. One cleavage site was between the two hydroxyl groups of the aliphatic chain (C30/C32, Fig. 2.B), second was between beta and gamma carbon of the β-OH-Asp on position 4 (C12/C13), corresponding to photolytic decarboxylation of this residue, and finally the third one between Gly and β-OH-Asp at the peptide backbone (C11/N).

Of the identified photolysis products, the most interesting was photolytic fragment PF1 with *m/z* 784.2420. A complementary HPLC analysis that preserves iron siderophore complexes revealed ion at *m/z* 837.1538. The observed *m/z* shows an isotope cluster compatible with the presence of 1 iron atom and corresponds to PF1 ([M+H]^+^; *m/z* 784.2420, error 4.3 ppm) with bound iron ([M-2H+Fe]^+^, 837.1538, error 3.7 ppm). According to MS/MS fragmentation, PF1 corresponds to cleavage of the aliphatic chain between carbons 30 and 32 and is accompanied by a reduction of the neighboring hydroxyl to a ketone (Fig. 2). With two β-OH-Asp residues present in the fragment, the fragment should retain its affinity to iron. It is unclear whether PF1 binds Fe^2+^ or Fe^3+^, but presence of PF3 (*m/z* 738.2367, error 4.4 ppm) suggests that PF1 (or eventually PF2) could undergo a second photolytic cleavage, upon which carboxyl group of β-OH-Asp on position 4 is cleaved of (C12/C13, see Fig. 2B).

### Effect of cyanochelin B on Synechocystis monoculture

To assess to what degree *Leptolyngbya* can monopolize iron by producing cyanochelin B, it was cultivated along with *Synechocystis* sp. PCC6803 as a reporter strain. *Synechocystis* does not produce its own siderophores and has growth rates and nutrient requirements similar to those of *Leptolyngbya*. First, we needed to verify that cyanochelin B has no additional inhibitory effect besides iron deprivation on *Synechocystis*.

The cultures of *Synechocystis* in standard and iron deprived media were exposed to cyanochelin B. Most importantly, the concentration of cyanochelin B had little to no effect on the culture grown in iron-deprived media. For example, at 0 µM and 180 µM final cyanochelin B concentration the optical density at 750 nm (OD) of *Synechocystis* increased from 0.2 to 0.65 and 0.6, respectively, indicating negligible inhibitory effect. The cultures grown in standard medium treated with cyanochelin B up to 2.2 µM reached approximately 1.5 times higher densities than those in iron-deprived media. In contrast, *Synechocystis* grown with cyanochelin B concentrations higher than 6 µM reached roughly the same density as their counterparts in iron deprived media. The results show that at a certain cyanochelin B:iron ratio the cyanochelin B can prevent *Synechocystis* from accessing iron. Furthermore, since the higher cyanochelin B concentrations do not produce additional growth reduction, the cyanochelin B has no additional inhibitory effect on *Synechocystis* other than the restriction of access. Cyanochelin B was dissolved in iron-deprived BG-11 and added to the test culture, yielding a final medium with approximately ∼10 µM of available Fe. However, in BG-11, iron tends to precipitate, and thus the actual concentration of biologically available iron may be lower and thus titrated already by 6.7 µM of cyanochelin B.

### Effect of UV-light on iron acquisition during cultivation

A co-culture plate assay was performed to test the influence of the photolytic activity of cyanochelin B on iron uptake and monopolization by *Leptolyngbya* sp. NIES 3755. *Synechocystis* cannot access the siderophore-bound iron directly (Fig. 3). However, it can take up the iron that is released by photolysis of cyanochelin-iron complexes and benefit from *Leptolyngbya*-produced cyanochelins indirectly. To test this, an experimental condition treated with UV radiation (and non-UV as control) was incorporated in this experiment. The combinatorial setup included three iron conditions: cultures deprived of iron (iron-deprived), iron provided as soluble ferric ammonium citrate (∼20 µM Fe, standard BG-11), and siderophore-accessible ferric chloride precipitates enclosed in alginate beads (bead-immobilized iron). Ferric ions were encapsulated in alginate beads to create an immobilized iron source. This locked form of iron should preferentially allow only siderophore-producers to access the iron, as low molecular weight siderophores can penetrate the alginate matrix and chelate the iron. This siderophore-iron complex would be available to siderophore producer strains or organisms with matching siderophore importer systems. For the experiment, 6-well plates with inserts, that provide 2 membrane (<1 µm) separated compartments with approximately the same volume (3 ml), were employed. This setting allows diffusion of compounds, prevents mixing of the model organisms and allows for their separate OD evaluation. Each compartment was inoculated with one of the experimental strains at OD of 0.02.

**Fig. 3.**
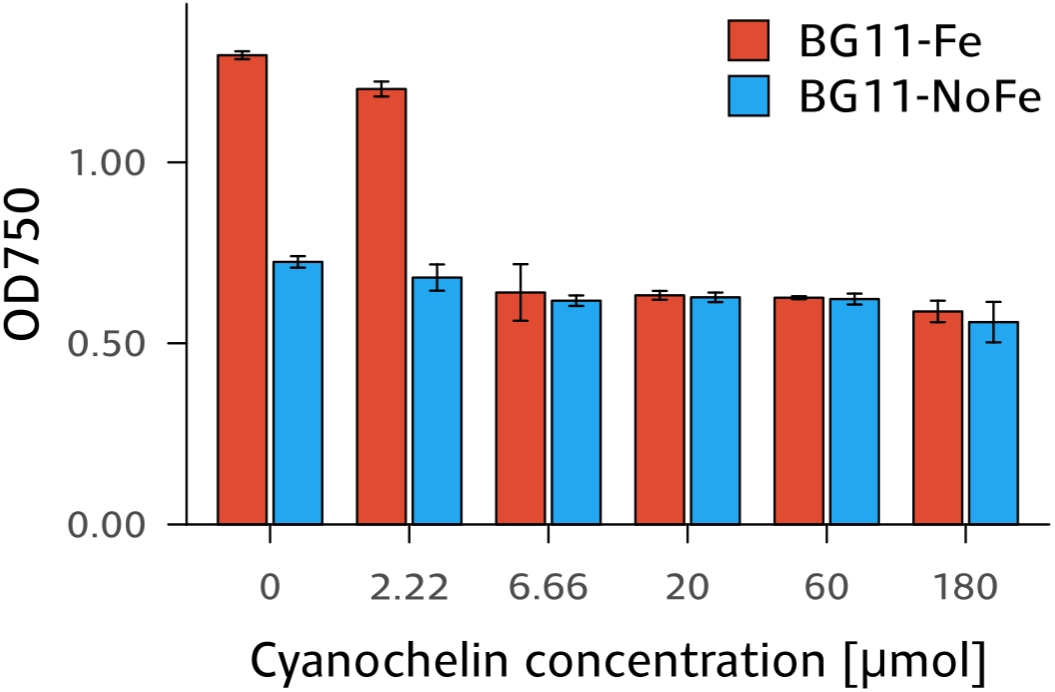
Endpoint OD of *Synechocystis* grown in presence of cyanochelin B in standard BG-11 and its iron-deprived version

In monoculture, *Leptolyngbya* grew well in the standard BG-11 and after 14 days of incubation it reached OD 1.48 and 1.64 in the non-UV and the UV conditions, respectively (Fig. 4). In presence of a bead-immobilized source of iron, the OD of *Leptolyngyba* culture increased to 0.89 (60% of the standard BG-11) and to 1.05 (64% of the standard BG-11) for non-UV and UV, respectively. This suggests that the producer takes up the siderophore-bound iron and iron released during photolysis with comparable/equal efficiency, if there is no other competing strain. In iron-deprived conditions *Leptolyngbya* reached OD 0.54 and 0.71 (36% and 43% of standard BG-11) for non-UV and UV conditions, respectively. Interestingly, in all 3 conditions *Leptolyngbya* grew slightly better in the presence of UV light, although this notion lacks proper statistical support.

**Fig. 4.**
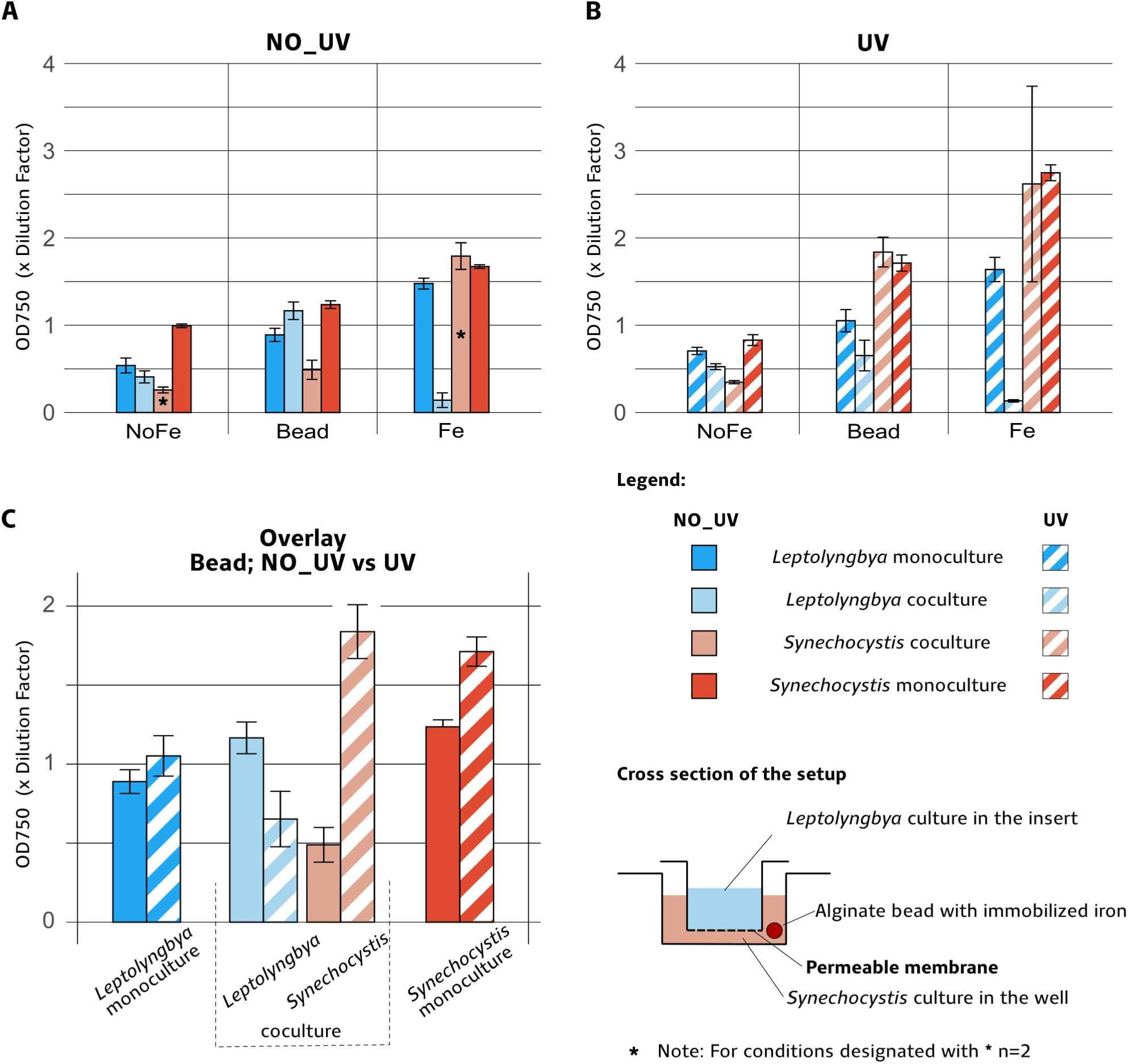
Cultivation experiments showing the combined effects of UV-light and iron availability on culture density of *Leptolyngbya* and *Synechocystis* cultivated individually or combined in membrane separated compartments. Endpoint OD of *Leptolyngbya* and *Synechocystis* grown under different availability of iron: iron deprived media - NoFe; alginate-enclosed iron - Bead; standard BG-11 medium (Fe) in the absence of UV - **A)** or presence of UV - **B)**. Subfigure **C)** shows comparison of endpoint ODs of cultures grown with alginate-enclosed iron in the absence or presence of the UV-light.

Compared to *Leptolyngbya*, monocultures of *Synechocystis* exhibited better growth in standard BG-11 and UV (OD 1.67 and 2.74 in the non-UV and the UV conditions, respectively, Fig. 4). In case of a bead-immobilized iron source, the final OD was 1.24 and 1.71 for non-UV and UV conditions, respectively, accounting for 74% and 62% of the ODs observed in standard BG-11. Similarly, in iron deprived conditions, *Synechocystis* reached ODs 0.99 and 0.82 in non-UV and UV conditions (59% and 30% of standard BG-11). Despite pre-starvation, *Synechocystis* managed to divide approximately 5 times (starting at OD 0.02), indicating that it either retained some reserves or was able to scavenge iron from the media. Also, *Synechocystis* could benefit from the immobilized iron bead to an extent comparable to *Leptolyngbya*. Complementary inductively coupled plasma mass spectrometry (ICP/MS) measurements estimated that an average alginate bead contains 2.08±0.21 µg Fe (mean±SD, n=9) and that our iron-deprived medium contained 1.43±0.23 µg/L (mean±SD, n=3), which corresponds to 700-times lower Fe content in comparison to standard BG-11 (∼1mg). Importantly, 3 mL of iron-deprived BG-11 incubated with 3 alginate beads for 13 days contained 12.47±0.12 ug/L Fe (mean±SD, n=3), accounting for less than 35 ng of iron. The amount of the leaked Fe is negligible copared to the amount of iron in the bead (∼0.017%), however, it is still ∼10x more free iron than what is available in the iron-deprived media.

In coculture, in standard BG-11, where iron is in excess and there is no competition for it, *Synechocystis* outcompeted *Leptolyngbya*. *Synechocystis* reached OD 1.79 and 2.62 in non-UV and UV conditions which accounts for 107% (non-UV) and 95% (UV) of corresponding monocultures. Interestingly, *Leptolyngbya* reached OD values of only 0.14 and 0.13 in non-UV and UV respectively, which corresponds to 12% and 8% of the OD values (0.539, 0.705) obtained in monoculture in the absence of iron. Perhaps, the limitation by other elements such as phosphorus or CO_2_ may hinder *Leptolyngbya* in coculture and make *Synechocystis* the better competitor.

The most interesting results were expected in the coculture with immobilized iron. In non-UV conditions *Leptolyngbya* reached OD 1.17 (71% of standard BG-11 and 125% of bead-immobilized iron *Leptolyngbya* monocultures) while cohabiting *Synechocystis* reached only 0.49 (27% OD in standard BG-11 and 40% of bead-immobilized iron monocultures). *Leptolyngbya* thus reached higher ODs than *Synechocystis* (∼2:1), which is in contrast with the cocultures in standard BG-11 medium and strongly suggests that under these conditions *Leptolyngbya* can efficiently monopolize the iron source. The presence of UV, however, cancelled the advantage and *Leptolyngbya* only reached OD 0.65 (56% of corresponding non-UV coculture, 40% of standard BG-11 UV and 62% of *Leptolyngbya* UV monoculture with bead-immobilized iron), while *Synechocystis* reached OD 1.84 which is 3.75 times more than in the corresponding conditions without UV and 117% of the OD of *Synechocystis* grown with *Leptolyngbya* in standard BG-11 and in presence of UV. At the end of the experiment, *Synechocystis* reached a ratio of 3:1 compared to *Leptolyngbya.* This result clearly indicates that the extent of monopolization is reduced under UV-light. Furthermore, it suggests that cyanochelin B releases free iron and supports the non-producer strain present in its vicinity.

The final ODs of *Leptolyngbya* in iron-deprived co-cultures reached 76% and 74% of their corresponding monocultures. For *Synechocystis,* the final ODs represent 26% and 42% of the reference monocultures. In iron deprived conditions, there is not much iron to compete for and is most likely taken up in the early phase of the experiment before the accumulation of critical amounts of siderophore in the media. Hence, there are no iron-cyanochelin complexes to photolyze and the effect of the UV on redistribution of iron via photolysis is not expected.

Even though there are many factors that influence the interaction between these two organisms, it is evident that when they are in close proximity to each other, photolysis of cyanochelin can change the dynamics of iron uptake and facilitates the growth of a non-producer by reducing iron monopolization.

### Phylogenetic distribution and biosynthetic gene clusters of cyanochelin B producers

During a field survey aimed at sampling of potentially iron-limited cyanobacterial communities that could produce novel siderophores, we obtained material that, after cultivation in iron-deprived media, contained cyanochelin B. Environmental sample number 146 was found as a microbial mat growing on a small wooden bridge within the splash zone of a waterfall in a calcareous region of inland Croatia (Fig. S18). Several cyanobacterial morphotaxa were identified in the original sample using light microscopy: *Microcoleus* sp., *Coleofasciculus* sp., *Calothrix* sp., and several Leptolyngbyaceae morphotypes. The sample was brought to the laboratory and kept in iron-depleted medium for approximately three months until the production of siderophores was induced and the LCMS analysis confirmed the presence of cyanochelin B. During cultivation and iron starvation, four *Leptolyngbya*-like morphotypes prevailed in the batch culture, of which single filaments were picked to isolate clonal strains (Table S4). PCR amplification of three different custom-designed *loci* (Table S5) across the cyanochelin B BGC was used to identify tentative producer strains. Three PCR-positive strains exhibited an identical morphology corresponding to *Phormidesmis* sp. (Fig. 5), therefore two additional strains of the morphotype were further examined. These isolates were cultivated in iron-depleted media and the production of cyanochelin B was confirmed by LCMS in four of them (S146-12, S146-33, S146-35, S146-36).

**Fig. 5.**
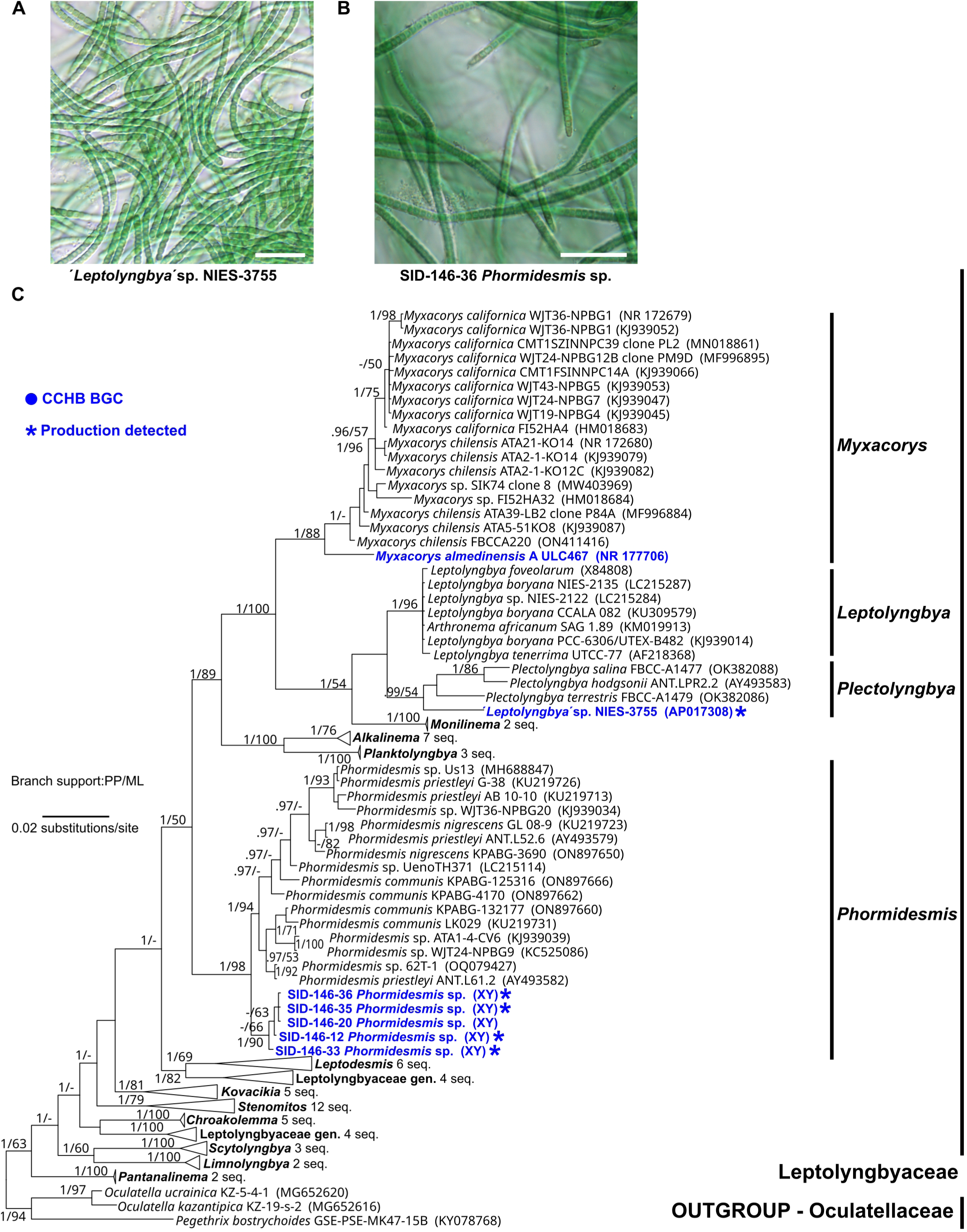
Morphology and phylogeny of cyanochelin B producers. **A)** and **B)** Microscopic view of the strains investigated in present study. C) Phylogenetic tree of 16S rRNA gene sequences featuring Leptolyngbyaceae taxa hosting biosynthetic gene clusters for cyanochelin B in their genomes. The production of cyanochelin B which was detected by HPLC-MS is marked by an asterisk. The topology of the tree represents Bayesian inference analysis. Supports of the branches, posterior probability (PP) and nonparametric bootstrap (ML) are presented. Scale bars represent 20 μm.

Genomes of all five strains were sequenced and of these four were successfully assembled. The genome assembly of the strain S146-36 failed due to contamination. The sequenced strains were searched for BGCs homologous to the cyanochelin B BGC as found in *Leptolyngbya* sp. NIES-3755. We confirmed the presence of cyanochelin B-compatible BGC in the three cyanochelin B producing isolates of *Phormidesmis* isolates, S146-12, S146-33, and S146-35 (Fig. 6.), but also in the strain S146-20, in which we could not confirm the production using HPLC-HRMS.

**Fig. 6.**
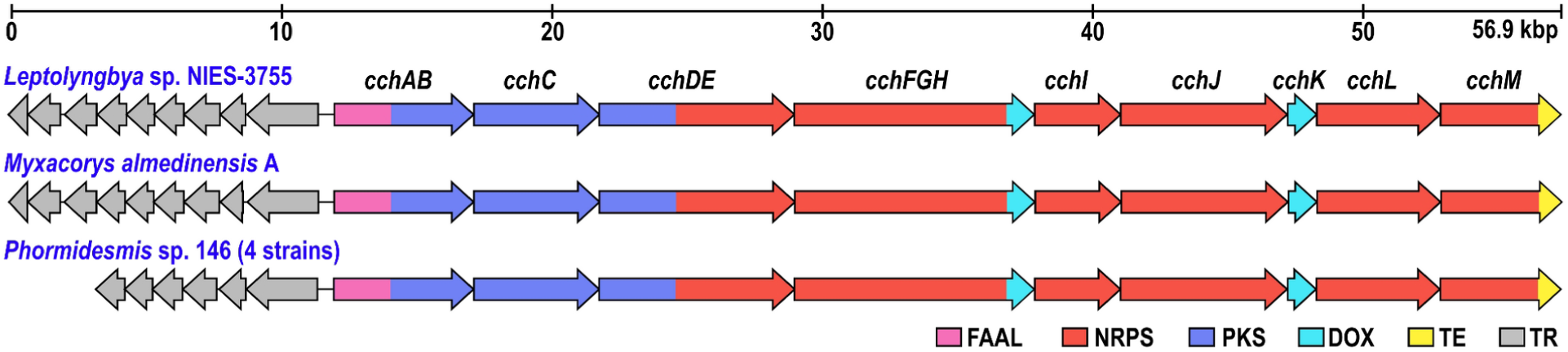
Arrangement of the cyanochelin B biosynthetic gene clusters. (BGCs) in strains of three separate genera of Leptolyngbyaceae (AP017310.1, NZ_WVIE01000017.1, sequences submitted to genbank, accession numbers to be retrieved soon). The NRPS/PKS biosynthetic core of all sequenced BGCs exhibits an identical topology and high average sequence identity (96%). Genes encoding siderophore transporters are found in opposite orientation adjacent to the first biosynthetic gene. FAAL - fatty acyl-AMP ligase; NRPS - non-ribosomal peptide synthetase; PKS - polyketide synthase; DOX - aspartate oxygenase; TE - thioesterase; TR - siderophore transport genes.

An additional BLAST search against the NCBI database identified a BGC highly similar to that of *Leptolyngbya* sp. NIES-3755 and *Myxacorys almedinensis* strain A (ULC467) (Fig. 6). Analysis of NRPS adenylation domains (A-domains) (Table S2) indicated that this strain is likely a cyanochelin B producer, however, its production capability was not investigated in the current study. The NRPS/PKS biosynthetic core of all analyzed BGCs was organized identically and consisted of nine genes spanning approximately 45 kbp, ordered co-linearly with the predicted biosynthesis (Fig. S2). Details of the BGC organization and encoded NRPS/PKS domains, including the predicted substrate specificity of A-domains, were previously described by Galica et al. (2021) for *Leptolyngbya* sp. NIES-3755, and are complemented here for *Phormidesmis* sp. 146 isolates and *M. almedinensis* A (Fig. S2, Table S2).

All examined *Phormidesmis* sp. strainś rRNA operon genes were almost identical, diverging only 0 - 0.3% in the 16S RNA gene and 0 - 0.7% in the ITS sequences, suggesting that all are representatives of a single species (Pecundo et al 2023). The cyanochelin B BGCs from the novel strains of *Phormidesmis* sp. represented three different genotypes with an average pairwise nucleotide identity of about 96% across the BGC. The sequence identity between the homologous proteins encoded in the BGC ranged from 93 to 100%.

The phylogenetic analysis of the 16S rRNA gene sequences from all three cyanochelin B-producing taxa and representatives of Leptolyngbyaceae did not support the existence of a separate clade with cyanochelin B-producers. While all of them were resolved as members of a single family, each of the three taxa harboring the BGC belonged to different genera, separated by a relatively long evolutionary distance (Fig. 5). The average pairwise nucleotide identity across the BGCs was slightly higher between *Leptolyngbya* sp. NIES-3755 and *M. almedinensis* A (90.6%), than between these two strains and *Phormidesmis* sp. (86.9 - 88.9%), which was also reflected in their average protein sequence identities (91.3% versus 87.7 - 89.4%). The difference was most prominent in the adjacent siderophore transporter gene cassette between *Phormidesmis* and the other two taxa (Fig. 6). In comparison to the set of nine genes presumably involved in cyanochelin B export/import in *Leptolyngbya* sp. NIES-3755 (Galica et al. 2021) and *M. almedinensis* A (Soares et al 2020), the *Phormidesmis* sp. isolates lacked three terminal genes, identified as homologues to DevA, DevB and DevC that encode an ABC transport system to possibly export the siderophore from cell interior to periplasmic space. However, a major facilitator protein (cctC - BAU16019) that is also possibly performing the same function is present in all the strains and these three genes may just code a functionally redundant system.

## Discussion

Microbial communities worldwide experience iron deprivation and under such conditions, siderophores can play a crucial role in determining the flow of biologically available iron. A substantial body of literature and experimental data show that the exclusive monopolization of iron by siderophore-producing organisms is not the prevalent scenario in complex microbial consortia. Instead, various forms of siderophore piracy and dependencies will likely emerge (Morris et al 2012, D’Onofrio et al 2010). The photolytic properties of certain siderophores introduce an additional layer of complexity, since such siderophores can support specific organisms dependent on access to Fe^2+^ that are present in the producer’s vicinity (Butler et al 2021, Barbeau et al 2001).

Siderophores are thought to be exchanged for dissolved organic carbon in mutualistic partnership of algae and bacteria (Barbeau et al 2001, Jiang et al 2024). However, cyanobacteria are versatile organisms that house both the capability to perform photosynthesis and generate fixed organic carbon from light and CO_2_ as well as the capability to produce siderophores and gain access to additional sources of iron. In this context, exploring which organisms are supported by cyanobacterial siderophores and what cyanobacteria could get in return is interesting. In the present paper we report our first steps to establish an experimental model to explore siderophore-mediated interactions between cyanobacteria and associated bacteria.

Our experimental setup employs particulate iron immobilized in alginate beads, membrane separation of the strains and lytic UV-A-light. We assume that in the environment iron is often not solubilized but present within a range of centimeters in some particulate form. By enclosing particulate iron in alginate beads we tried to replicate such conditions. The setup also included a chemically inert membrane to separate the strains under study while allowing diffusion. Importantly, the bead was placed in the compartment with the siderophore non-producing strain to ensure the siderophore would pass the membrane on its way to the iron source. Microbial communities often exhibit complex spatial organization with overlapping distribution of individual strains across multiple gradients (Whitton 2012). The lack of spatial organization should not considerably affect our observation of siderophore routing of iron. In the experiments, we have used constant light composed of warm-white light and UV-A provided by LED strips. Reproducing natural light with all its components and variations in a lab is impossible. However, it was important to include a light that would cause photolysis of cyanochelin B, that we discuss below.

Cyanochelin B is an amphiphilic lipo-heptapeptide that employs two β-OH-Asp residues and likely a combination of hydroxyls and a thiazoline ring for iron chelation. This unique structure contributes to the repertoire of known siderophores from cyanobacteria. β-OH-Asp is broadly used as an iron chelating residue and is found in 124 of 707 (17.5%) siderophores cataloged in a recently updated siderite database (He et al 2024). The frequent occurrence of β-OH-Asp in siderophores suggests that photolysis of siderophores may be a widespread and currently underestimated phenomenon in microbial ecology. In cyanochelin B, the two β-OH-Asp residues account for four out of six electron pairs required for chelation of ferric iron. Currently we lack experimental data to unambiguously determine the origin of the remaining two electron pairs. However, on the N-terminal moiety of the peptide, there is a thiazoline linked to a hydroxylated acyl chain. The relative position of the nitrogen in the thiazoline and hydroxyl group on the α-carbon of the acyl chain may provide the required electron pairs, similarly as hydroxyphenylthiazoline found in yersiniabactin or alpha-hydroxyimidazole present in corrugatin (Miller et al 2006, Mular et al 2024, Risse et al 2014). Furthermore, photolytic cleavage typically occurs near the iron-binding residues (Hardy and Butler 2019) hence the observed cleavage at C30-C32 suggests that at least one of the hydroxyls in its proximity is in contact with iron. Confirming the role of these hydroxyls in iron chelation was beyond the scope of this study, however, if it was confirmed by NMR data it would be a novel structural motif employed for iron chelation.

Photolysis of cyanochelin produces several types of fragments. Their relative abundance cannot be accurately determined due to possibly different ionization at the MS source and lack of analytical standards for the individual compounds. However, assuming the same efficiency of MS-ionization for all the fragments, it seems that PF1 and PF2 are the most abundant. Interestingly, at least one of the fragments, particularly PF1, seems to retain the ability to chelate iron, similarly as was reported for pacifibactin, petrobactin, aerobactin and aquachelin (Hardy & Butler 2019, Butler 2021, Kreutzer et al 2012). Less explored is the possible capability of a photolytic fragment to perform an additional round of photolysis. Our data suggest that PF1 can undergo a secondary photolysis. However, the fragment must be isolated and tested separately to properly prove it. Although such capability was considered for e.g. aquachelin, we are currently unaware of any described case.

Cyanochelin B reduces ferric iron to its ferrous form. The measured rates and half-lives of the iron-siderophore complex (maximal rate 0.078 min^−1^ or 4.7 h^−1^; t_1/2_ ∼9 min) suggest that once iron is bound by the siderophore it immediately undergoes photoreduction within the range of minutes. Amin and colleagues reported photolytic rates of vibrioferrin (0.031 h^−1^), petrobactin (0.003 h^−1^) and marinobactin (>0.001 h^−1^) illuminated by fluorescent light at intensity 80 µE m^−2^ s^−1^ (Amin et al 2009). The authors also measured photolysis of vibrioferrin under attenuated sunlight at intensity of 500 µE m^−2^ s^−1^ to later multiply it by 4 (12.9 h^−1^, t_1/2_ ∼3.2 min) in order to compare it with previously reported rates of photolysis of aquachelin under sunlight in 2000 µE m^−2^ s^−1^ (0.6 h^−1^). Comparing the rates of photolysis of individual siderophores is complicated. First, for the structural differences: petrobactin and vibrioferrin both use a citrate moiety, however the former at a central position and the later at a terminal position; and marinobactin employs β-OH-Asp that is further cyclized to a 9-membered heterocycle while aquachelin uses a simple β-OH-Asp. These differences affect the absorbance spectra as well as the possible course of reaction. Second, the spectra of the used light-source are not usually available, hence in combination with different absorption spectra of the siderophores it is not possible to clearly state how much of the photolytic light is being applied. For UV-dependent photolysis we have used UV-A emitting LED lights that have an emission peak at 360 nm at intensity ∼3.5 µE over the range 315-415 nm that we also used for our cultivation experiments. Even at this intensity the rates for cyanochelin B are higher than for aquachelin in direct sunlight and 36% of the calculated values for vibrioferrin. Overall, it can be concluded that cyanochelin undergoes photolysis at relatively high rate even at intensity of UV light that corresponds to light conditions of a cloudy day in temperate region. In cultivation with bead-enclosed iron, the rate of photolysis is also influenced by the kinetics of formation of iron-siderophore complexes. In nature, the process is likely further complicated by the changing light conditions and much more complex chemical milieu that often includes UV-protective pigments (Whitton 2012). Also, we have used constant light, and did not include “night” in our cultivation experiments, during which the pool of cyanochelins could possibly replenish and direct the flow of iron to a different set of microorganisms within the community.

Siderophores are frequently considered to be involved in complex microbial interactions (Kramer et al 2019). The possible role of photolytic siderophores in microbial communities has been considered previously, however, in the context of relationship between phototrophic algae and siderophore-producing bacteria (Barbeau et al 2001). In such case, the exchange of fixed organic carbon for siderophore-mediated access to additional iron sources seems to be a mutually beneficial trade. In our experimental model we employ two phototrophic organisms that compete for the same set of nutrients and of which only one can produce siderophores. *Synechocystis*, the non-producer, outcompetes the siderophore-producing *Leptolyngbya* in standard cultivation media. Such a situation is unlikely to occur in the alga-bacteria system, where bacteria live on algal exudates. When our model strains are set to compete for a limited source of iron, *Leptolyngbya* is clearly able to monopolize access to it in the absence of UV-light. In the presence of UV, however, the photoreduced iron is also available to *Synechocystis* in amounts sufficient to regain its advantage. In the presence of UV light, *Leptolyngbya* clearly supports its direct competitor and is likely to become an indispensable minority in the model system. This is in accordance with the postulates of the Black Queen Hypothesis, which expects the occurrence of strains that form a minority but are retained since they carry the load of an essential metabolic capability crucial for the survival of the whole community (Morris et al 2012).

Cyanochelin B was detected in iron-starved cultures of *Leptolyngbya* sp. NIES-3755 (Hirose et al 2016) and in four strains of *Phormidesmis* described in the current study. In addition, cyanochelin B BGCs were found in one more strain of *Phormidesmis* and in *Myxacorys almediensis*. All of the concerned strains are simple filamentous cyanobacteria from the Leptolyngbyaceae family (*sensu* Mai et al 2018, Strunecký et al 2023). They are found worldwide to grow subaerophytically on exposed soil (*Leptolyngbya,* Hirose et al 2016) but also on man-made structures, such as stony walls (*Myxacorys,* Soares et al 2020) and wooden bridges (*Phormidesmis,* current study). Subaerophytic microbial mats, however, often lack a reliable access to required nutrients and face severe fluctuations of light intensity, temperature and water availability. Such mats are frequently found and are ecologically important in extreme environments, such as deserts, high altitude mountains and polar regions. It is tough to address the availability of iron for microbial mats. Frequent changes of environmental conditions and possibly steep gradients of pH and redox potential within the microbial mats make it almost impossible to estimate when and whether a particular filament is iron-limited. However, it is reasonable to estimate the cyanobacteria in microbial mats will undergo at least intermittent periods of iron deprivation. In the exposed microbial mats, filamentous cyanobacteria, such as *Leptolyngbya* or *Phormidesmis,* are frequently the keystone species. They provide fixed carbon, possibly also nitrogen, and their filamentous bodies are important structural features that provide shelter for a multitude of associated microbes (Oren et al 2017, Campbell et al 1979). In our hands, however, the cyanochelin-producing species was not dominant and only rose to prominence during prolonged cultivation in iron-deprived medium and in the absence of UV light.

The strains with cyanochelin B-encoding BGC all belong to a sub-lineage of *Leptolyngbyaceae* consisting of the genera *Leptolyngbya*, *Plectolyngbya*, *Myxacorys*, *Monilinema*, *Alkalinema*, *Planktolyngbya* and *Phormidesmis* (Strunecký et al 2023). The cyanochelin BGC organization (Fig. 6) and sequence identity patterns between the individual biosynthetic genes correspond with the reconstructed evolutionary distances between the three taxa (Fig. 5). The metabolic capability to synthesize cyanochelin B could be an ancestral trait although that would require a high number of gene loss events during which the BGC is compromised. The Black Queen Hypothesis postulates that such events may be frequent, if there is another member of the microbial community to compensate for the lost metabolic capability (Morris et al 2012, Tostado-Islas et al 2021, D’Onofrio et al 2010). The concerned phylogenetic lineage, however, includes multiple other strains, i.e. *Leptolyngbya boryana* NIES-2135, *Myxacorys chilensis* and *Myxacorys californica,* that harbor BGCs that could possibly encode biosynthesis of cyanochelin-like siderophores with 2 β-OH-Asp included in their structures (Strunecký et al 2023, Galica et al 2023). The situation around evolution of cyanochelin B BGC is further complicated by the fact that in *Leptolyngbya* sp. NIES-3755 the cyanochelin BGC is located on a plasmid (Galica et al. 2021) and can thus easily undergo horizontal gene transfer.

Considering the frequency of β-OH-Asp found in siderophores, the photolysis of iron-siderophore complexes and accompanied release of reduced iron may be a phenomenon with underestimated effect on cycling of iron in microbial communities and potentially on global Fe cycle. Cyanochelin B is a prominent example of such siderophores that is found in the cosmopolitan Leptolyngbyaceae family on which we show that the redistribution of photoreduced iron may occur without specific targeting and can altruistically support any microbe, including direct competitors of the producer.

## Methods

### Strains and cultivation

*Leptolyngbya* sp. strain NIES-3755 was obtained from NIES culture collection and grown on BG-11 medium at 21°C and continuous dispersed light. For production of the compound, the strain underwent several passages in iron depleted medium until siderophore production was confirmed by HPLC-HRMS and then harvested. *Synechocystis* sp. PCC 6803 Nixon was kindly provided by prof. Roman Sobotka (Centre Algatech, Institute of Microbiology of the Czech Academy of Sciences) and kept at 28°C.

Iron-deprived media was prepared analogously to standard BG-11 but without the addition of ferric ammonium citrate. The pH of BG11 media was adjusted to 7.5 (Stanier et al 1971).

### Siderophore inhibition test

*Synechocystis* was grown for ∼7 days in iron deprived media at 25°C and 50 µE with constant bubbling until it reached OD750 = 0.8 (1cm optical path, V-1200 Spectrophotometer, VWR). The culture was spun down by centrifugation (3000 rpm for 5 minutes, benchtop centrifuge Eppendorf). The cell pellets were rinsed by ironless medium, pelleted again and finally resuspended in both standard and ironless media to final OD 750 = 0.4. The cell suspensions were used to inoculate a 96-well plate. The treatment solution consisting of ironless medium and cyanochelin B at various concentrations was added to the cell suspension in 1:1 ratio yielding the cultures at starting OD750 = 0.2 and cyanochelin B concentrations of 180, 60, 20, 6.7, 2.2 and 0 µM. The experiment was evaluated after 10 days of cultivation at 50 µE and 25°C by measuring OD750 by Omega Fluostar plate reader.

### Preparation of alginate beads

To prepare alginate-enclosed precipitate iron, an aqueous solution of ferric chloride (0.5M) was precipitated by addition of 1M NaOH in 1:9 ratio and pelleted; the supernatant was removed, and the pelleted iron precipitates were mixed with aqueous solution of sodium alginate to final 1% (w/w) alginate. The suspension was dropped into a continuously-stirred solution of calcium chloride (0.1M) to produce ∼ 40 µL beads, each containing roughly 2.15 µg Fe.

### Iron determination

Determination of iron in alginate beads/scads and cultivation media was performed by inductively coupled plasma mass spectrometry (ICP-MS). An Agilent 8900 ICP-MS/MS instrument equipped with MicroMist concentric nebulizer, Scott double-pass spray chamber, quartz torch with 2.5 mm i.d. injector and nickel sampling/skimmer cones was used for quantification of iron in digested (beads and scads) or diluted (media) material. ICP-MS/MS was operated at 1550 W input power, 8 mm sampling depth and 1.20 L/min flow of carrier gas (Ar). Octapole reaction system operated in collision mode with helium flow 10 ml/min was used for suppression of spectral interferences.

Prior analysis, alginate beads and scads were mineralized with 1 ml of concentrated nitric acid (Merck, suprapure) and 0.5 ml of hydrogen peroxide (Merck, empsure). Samples were digested at 170 °C for 20 minutes with ramping of temperature in 20 minutes using microwave decomposition system (MARS 6, CEM Corporation). Finally, mineralized samples were diluted to 10 ml with Milli-Q type 1 ultrapure water (Milli-Q Direct, Milliore). Cultivation media were 5x diluted to 1% nitric acid prior analysis.

### Co-cultivation experiments

*Leptolyngbya* and *Synechocystis* were cultivated individually or in membrane-separated compartments (ThinCerts®, Greiner Item-No. 657641). The cultures were either deprived of iron, or the iron was provided as soluble ferric ammonium citrate (∼20 µM Fe, standard BG-11) or as ferric chloride precipitates enclosed in alginate beads, referred to as bead-immobilized iron, which is supposedly accessible by siderophores.

The cultures were first primed in full iron BG11 medium followed by a period of iron starvation for a week. Pre-starved cultures of cyanobacteria were inoculated to the 6-well plates at an initial OD 750 = 0.05 (1cm optical path, V-1200 Spectrophotometer, VWR). *Leptolyngbya* was inoculated into the insert compartment, and *Synechocystis* was placed below the insert into the well. The inserts were also added to monocultures to avoid any shading bias. Plates were placed in UV (1.9 µE, UV-A, 350-390 nm) and non-UV conditions to determine the effect of cyanochelin B photolysis. The plates were incubated for 15 to 20 days at 25°C with shaking and continuous light (warm white LED) at 50 µE. At the end of the experiment, the cultures were thoroughly mixed and their OD750 was measured on V-1200 Spectrophotometer (VWR) in a 1 ml disposable plastic cuvette with a 1 cm optical path.

### HPLC-HRMS analysis

The concentration of cyanochelin B and its photolytic fragments was analyzed on Dionex Ultimate 3000 HPLC system coupled to Bruker Impact HD II mass spectrometer equipped with electrospray ionization. Chromatographic separation was achieved on Waters MaxPeak XBridge Premier BEH C18, 130 Å, 2.5 μm; 50×2.1 mm column. Acetonitrile (A) and water (B), supplemented with trifluoroacetic acid at concentrations 0.002% and 0.1% v/v were used as mobile phases. The column was eluted with the following gradient- 0 min 5/95%; 2 min 5/95%, 8 min 100/0%, 10 min 100/0%, 11 min 5/95%, and 12 min 5/95%. at a flow-rate of 0.4 ml/min. The proportion of the organic mobile phase between 2 and 8 minutes was increased non-linearly by convex upward curve 2 available in Chromeleon. The electrospray ion source dry temperature was set to 250°C, drying gas flow to 11 liters/min and nebulizer pressure to 3 bars. Capillary voltage was set to 4,500 V and endplate offset to 500 V. The spectra were collected in the range of 50 to 1,490 *m/z* with a spectral rate of 2 Hz. Precursor ions were selected automatically and collision-induced dissociation (CID) was set as a ramp from 20 to 50 eV. Calibration of the instrument was performed using CH_3_COONa clusters at the beginning of each analysis. Data were analyzed using Bruker Data Analysis software and MZmine3 (Schmidt et al 2023).

### Isolation of cyanochelin B

Spent media of iron-deprived culture of *Leptolyngbya* with HPLC-HRMS-confirmed content of cyanochelin B was shaken overnight with amberlite XAD-16, then rinsed with 10% MeOH and eluted with 100% MeOH. The extracts were evaporated on a rotary evaporator (70mBar, 40°C) and stored in the freezer. Before HPLC separation, the extracts were dissolved in a minimal volume of MeOH. The compounds were isolated on a µBONDAPAK phenyl column (7.8mm X 30mm, WATERS) using acetonitrile (A) and 25% v/v acetonitrile-water mixture (B). The solvents were amended with TFA (0.002% and 0.08% v/v, respectively), which proved to be crucial for preventing unspecific compound-column interactions and ensuring reliable elution. The column was eluted with the following linear gradient: 0 min, 100% B, 3 ml/min; 2 min, 100% B, 3 ml/min; 4 min, 100% B, 4 ml/min; 8 min, 50% B, 4 ml/min; 24 min 65% B, 4 m;/min; 25 min, 100% B, 4 ml/min; 28 min, 100% B, 4 ml/min; 29 min, 0%B, 3 ml/min; 30 min, 0% B, 3 ml/min. Column temperature was maintained at 40°C. Fractions were collected by an automated fraction collector, on a time basis with 0.75-minute windows starting from minute 2 and ending at minute 27. Collected fractions were checked for the content of the compound of interest by HPLC-HRMS (see above), pooled accordingly and dried. If needed, pooled high 1026-content prepurified extracts were subjected to a second round of HPLC isolation on the same column with an analogous MeOH/water gradient. The obtained fractions were selected based on the data and evaporated.

### Structural characterization of cyanochelin B

^1^H NMR and 2D NMR experiments were carried out at 35°C on a Bruker AvanceNeo 700 MHz spectrometer (Billerica, MA, US) equipped with a triple resonance CHN cryoprobe, using d-DMSO (Sigma Aldrich, Milan, Italy); chemical shifts were referenced to the residual solvent signal (d-DMSO: δ _H_ 2.50 ppm, δ_C_ 39.51 ppm.). The HSQC spectra were optimized for ^1^J_CH_ = 145 Hz. The HMBC experiments for ^2,3^J_CH_ = 8 Hz. Abbreviations for signal couplings are as follows: s = singlet, d = doublet, br.d = broad doublet, dd = doublet of doublets, td = triplet of doublets, t = triplet, q = quartet, m = multiplet.

### Advanced Marfey’s analysis

The cyanochelin B (10 μg) was hydrolyzed with 600 μL of 6N HCl/AcOH (1:1) at 120 °C for 18 h. The residual HCl fumes were removed under an N_2_ stream. The hydrolysate was then dissolved in TEA/acetone (2:3, 200 μL) and the solution was treated with 200 μL of 1% 1-fluoro-2,4-dinitrophenyl-5-L-alaninamide (L-FDAA) in CH_3_CN/acetone (1:2) (Esposito et al 2016). Subsequently, another aliquot of cyanochelin B (10 ugs) was hydrolyzed, as above, and derivatized with 200 μL of 1% 1-fluoro-2,4-dinitrophenyl-5-D-alaninamide (D-FDAA). The vial was heated at 50°C for 2 h. The mixture was dried, and the resulting L-DAA and D-DAA derivatives of the free amino acids were redissolved in MeOH (200 μL) for subsequent analysis. Authentic standards of L-Tyr, D-Ser and L-β-OH-Asp were treated with L-FDAA and D-FDAA as described above and yielded the L-DAA and D-DAA standards. Marfey’s derivatives of cyanochelin B were analyzed by using the high-performance liquid chromatography system (Thermo U3000 HPLC system) connected to a Thermo LTQ Orbitrap XL mass spectrometer (Thermo Fisher Scientific Inc.,Waltham, MA, USA), and their retention times were compared with those from the authentic standard derivatives. A 5 μm Kinetex C18 column (50 × 2.10 mm) maintained at 25 °C was eluted at 200 μL min^−1^ with 0.1% HCOOH in H_2_O and ACN. The gradient program was as follows: 5% ACN 3 min, 5–50% ACN over 30 min, 50-90% ACN over 1 min and 90% ACN 6 min.

Mass spectra were acquired in positive ion detection mode. MS parameters utilized a spray voltage of 4.8 kV, a capillary temperature of 285 °C, a sheath gas rate of 32 units N_2_ (ca. 230 mL/min), and an auxiliary gas rate of 15 units N_2_ (ca. 150 mL/min). The MS method involved four HRMS/MS scans after each full MS scan for the four most intense ions detected in the spectrum (data-dependent acquisition mode, DDA). The *m/z* range for data-dependent acquisition was set between 150 and 2000 amu with resolution set to 60,000. HRMS/MS scans were obtained for selected ions with CID fragmentation using an isolation width of 4.0, a normalized collision energy of 35, an activation Q of 0.250, and an activation time of 30 ms. Mass data were analyzed using the Xcalibur suite of programs.

Analysis of mass spectra of both β-OH-Asp standards and the cyanochelin-L-DAA and cyanochelin-D-DAA were carried out by using a 3 μm Phenomenex Luna Omega Polar C18 (100 × 2.10 mm) maintained at 45 °C and eluted at 400 μL min^−1^ with 0.1% HCOOH in H_2_O and ACN. The gradient program was as follows: 10% ACN 1,5 min, 10–95% ACN over 8,5 min and 95% ACN 2 min.

The analysis indicated that the amino acid forms various adducts with the derivatizing agent, prompting further analysis of chromatograms from LC-MS SRM (Selected Reaction Monitoring) spectra. In SRM, mass analyzers are set to a selected mass-to-charge ratio, focusing on specific precursor and product ions. The MS method involved two HRMS/MS scan events, following two product ions: *m/z* 356.1-358.1 and *m/z* 280.1-282.1 from the *m/z* precursor 402.2.

### Determination of kinetic of cyanochelin B

Cyanochelin B (10 µM) was mixed with excess FeCl_3_ (20 µM) in 10mM Ammonium acetate buffer (pH = 5) and incubated overnight at room temperature to ensure formation of iron-siderophore complexes. The buffer was chosen for its compatibility with MS. A control mixture without addition of FeCl _3_ was prepared analogously. Aliquots of the solution (75 µL) were distributed to individual round-bottom wells of a 96-wellplate (VWR art.no. 734-2803). The plate was placed under source of UV-A and visible light provided by LED strips. The intensity of visible light was constant in all variants of the experiment (50 µE). Four intensities of UV light were used: UV0 - UV-A light source was off; UV1 - UV-A 0.9 µE; UV2 - UV-A 1.9 µE; UV3 - UV-A 3.5 µE; and UV4 - UV-A light source set to 1.9 µE and the well plate was covered with the lid as during cultivation of cyanobacteria. At given time points the timer was stopped, the plate was removed from the source of UV light and 75 µL of MeOH was added to the well. The content of the well was mixed thoroughly and the solution was transferred to a glass vial. The plate was returned to designated light conditions and the timer was resumed. The sample was protected from light and the sample was analyzed by HPLC-HRMS within 24 hours. The data were processed using MZmine3 to quantify content of cyanochelin B. Linear regression was applied on the log transformed integrated peak areas of cyanochelin B to confirm the order of the reaction and determine its rate.

### Field sampling and strain isolation

A microbial mat growing on a splashed surface of a wooden bridge over a stream near a waterfall at GPS: N 44.04152, E O16.23488, altitude 87 m, was collected as sample 146 by scraping the surface with an ethanol-cleaned spatula and placed in a sterile 50 ml tube. Later the same day, the sample was examined by an Olympus CX23 light microscope equipped with an oil-less 60x objective. After confirmation that the mat was dominated by cyanobacteria, the sample was split into: a) disposable plastic cultivation bottles in iron-deprived and standard Z medium (Staub 1961); b) a sterile cryo-vial and placed in a travel container with liquid nitrogen to preserve the original community for DNA analysis; c) smashed and preserved with 20% glycerol and stored as a backup. The sample was then kept in a travel fridge (∼8°C) with a LED light to transfer the live community. Once the multispecies culture derived from sample 146 was confirmed to produce cyanochelin B by HPLC-MS analysis, it was inspected by optical microscope and single filaments of the dominant morphotypes were picked by a sterile glass capillary into fresh Z medium (Rippka 1988).

### Cyanochelin B PCR protocol

Based on sequences of the previously identified cyanochelin B BGC of *Leptolyngbya* sp. NIES-3755 (Galica et al 2021) and a homologous BGC harbored in the genome of *Myxacorys almedinensis* A (ULC467 - GCA_010091945.1) disclosed by the BLAST search, a PCR protocol amplifying five loci scattered among the cluster was designed (Table S5). Whole genomic DNA from sample 146 and from *Leptolyngbya* sp. NIES 3755 (applied as a positive control) was isolated using NucleoSpin Soil Mini kit (Macherey Nagel, Düren, Germany) following the manufacturer’s instructions. The PCR amplification was performed using each loci-specific primer pairs and sent for Sanger sequencing (SeqME, Dobříš, Czech Republic) as described in Štenclová et al (2023). The protocol confirmed the occurrence of cyanochelin B BGC in the mixed sample 146, being positive in three loci (1, 4 and 5). Moreover, the occurring single nucleotide polymorphisms indicated cluster variants. The designed protocol was further used to screen isolated clonal strains to identify the potential producers. DNA from three strains of each *Leptolyngbya*-like morphotype was isolated (Tab S4), PCR executed, and positive loci prepared for sequencing (as described previously). Taxonomic identity of three strains of all four morphotypes was confirmed by 16S rRNA PCR amplification and sequencing using VRF2 (5‘– GGG GAA TTT TCC GCA ATG GG –3‘) VRF1 (5‘– CTC TGT GTG CCT AGG TAT CC –3’) primers and PCR setting described in Johansen et al. (2011).

### Whole genome sequencing and assembly

Single filaments of each of the five clonal strains of *Phormidesmis* sp. from sample 146 were isolated by glass capillary technique and the genomic DNA was amplified using multiple-displacement amplification (MDA) with the Repli-G Mini Kit (Qiagen, Hilden, Germany) as described previously (Mareš et al. 2014). MDA products from 5–8 filaments per strain that passed a quality check by 16S rRNA sequencing, were pooled together and sent for de novo genome sequencing (Institute of Experimental Botany, Centre of Plant Structural and Functional Genomics, Olomouc, Czech Republic) using an Illumina HiSeq Pair-End library with 150 bp reads and 2.5 Gbp data yield per sample. The raw data from Illumina were trimmed, assembled, binned to remove minor bacterial contamination, and taxonomically classified using the *nf-core/mag* metagenomics pipeline (Krakau et al. 2022) installed on the server of the Institute of Hydrobiology, Biology Centre of the CAS.

### Phylogenetic analysis

The final rRNA operon data of *Phormidesmis* sp. SID-146-12, 20, 33 and 35 were excised from their sequenced genomes (Geneious Prime 2024.0.7), for *Phormidesmis* sp. SID-146-36 was obtained only a partial sequence from a sanger-sequenced PCR product amplified by the primers VRF2 (5‘– GGG GAA TTT TCC GCA ATG GG –3‘, Johansen et al. 2011) and 1494 (5‘– CTA CGG CTA CCT TGT TAC GA –3‘, Weisburg et al 1991). Data was deposited in the GenBank - NCBI database under the accession numbers PQ675769–PQ675773. Obtained sequences were aligned with a wide range of Leptolyngbyaceae representatives, including all major lneages inside the family. The final matrix consisted of 108 taxa, including 3 taxa of Oculatellaceae employed as an outgroup and it was 1112 positions long. The alignment was constructed using MEGA11 (Tamura et al. 2021) by the MUSCLE (Edgar 2004) algorithm and adjusted manually. Bayesian inference was executed in the program MrBayes v. 2.6 (Ronquist et al. 2012) with default settings for 5 million generations. Posterior probabilities of data split were recorded for individual branches. Maximum likelihood analysis was performed in the PhyML software (Guindon et al. 2010) applying the best model TN93+R automatically chosen by SMS algorithm (Lefort et al. 2017). Nonparametric bootstrap of 1000 pseudo replications was calculated and used as branch support. Obtained trees were merged and the final figure was adjusted in graphic illustrator InkScape (https://inkscape.org/). Only supports above 50 (ML) and above 0.95 (PP) are displayed. For novel strains, check of p-distances of the rRNA as well as cyanochelin B BGC similarity was done by MEGA11.

### Analysis of the cyanochelin BGCs

The genomic bins of *Phormidesmis* sp. strains were automatically screened for secondary metabolite BGCs using the antiSMASH v. 7.0 online analysis tool in bacterial version (Blin et al. 2021). Contigs containing tentative BGCs with similarity to cyanochelin A in KnownClusterBlast were analyzed in detail to address the composition of NRPS and PKS domains, the predicted substrate specificity of amino acid adenylation domains (Stachelhaus code), the presence of amino acid epimerase domains, and identity of additional tailoring enzymes to aid with cyanochelin structure elucidation and to link the BGC to its product. Stereoconfiguration of the PKS ketoreductase (KR) domain substrate was predicted based on an alignment of bacterial KR domains of various types together with the KR domain encoded in the *ccsAB* gene, using an analysis of conserved residues within the active site groove (Keatinge-Clay 2007).

## Acknowledgements

This research was partly supported by the National Biodiversity Future Center (NBFC), in the frame of European Union—NextGenerationEU and the European Marine Biological Resource Centre (EMBRC), project nr. 121 06/21/2022 (IR0000035)” Unlocking the Potential for health and by “Finanziamento della Ricerca di Ateneo” (FRA) 2022, by Università degli Studi di Napoli Federico II, with the contribution of Compagnia di San Paolo.

Authors thank the RECETOX Research Infrastructure (No LM2023069) financed by the Ministry of Education, Youth and Sports for supportive background.

